# Comparative single-cell transcriptomes of dose and time dependent epithelial-mesenchymal spectrums

**DOI:** 10.1101/2022.05.06.490972

**Authors:** Nicholas Panchy, Kazuhide Watanabe, Masataka Takahashi, Andrew Willems, Tian Hong

**Affiliations:** Department of Biochemistry and Cellular and Molecular Biology. The University of Tennessee, Knoxville. Knoxville, Tennessee, 37996, USA; RIKEN Center for Integrative Medical Sciences, 1-7-22 Suehiro-cho, Tsurumi-ku, Yokohama, Kanagawa 230-0045, Japan; School of Genome Science and Technology, The University of Tennessee, Knoxville. Knoxville, Tennessee 37916, USA; National Institute for Mathematical and Biological Synthesis, Knoxville, Tennessee, 37996, USA

**Keywords:** Single-cell RNA-sequencing, Epithelial-mesenchymal transition, Phenotypic plasticity, TGF-β, Dose-dependent cellular responses

## Abstract

Epithelial-mesenchymal transition (EMT) is a cellular process involved in development and disease progression. Intermediate EMT states were observed in tumors and fibrotic tissues, but previous *in vitro* studies focused on time-dependent responses with single doses of signals; it was unclear whether single-cell transcriptomes support stable intermediates observed in diseases. Here, we performed single-cell RNA-sequencing with human mammary epithelial cells treated with multiple doses of TGF-β. We found that dose-dependent EMT harbors multiple intermediate states at nearly steady state. Comparisons of dose- and time-dependent EMT transcriptomes revealed that the dose-dependent data enable higher sensitivity to detect genes associated EMT. We identified cell clusters unique to time-dependent EMT, reflecting cells *en route* to stable states. Combining dose- and time-dependent cell clusters gave rise to accurate prognosis for cancer patients. Our transcriptomic data and analyses uncover a stable EMT continuum at the single-cell resolution, and complementary information of two types of single-cell experiments.

## Introduction

Epithelial-mesenchymal transition (EMT) is a cellular process in which epithelial (E) cells undergo fate switches towards mesenchymal (M) types. This process renders the loss of apical-basal polarity and the gain of migratory properties. EMT plays crucial roles in development and disease progressions such as metastasis and fibrosis [1, 2]. EMT is not a binary process. In tumor cells, for example, intermediate (partial) EMT states were observed [3-5], and it was suggested that there is an association between intermediate EMT states and metastatic potentials [6, 7]. Interestingly, intermediate EMT states can also be observed *in vitro* with epithelial cell lines treated with EMT signals, such as TGF-β [3, 8], and these *in vitro* experiments provide useful insights into molecular programs underlying partial EMT [9]. For example, experiments with genetically perturbed cells have suggested that interconnected feedback loops in gene regulatory networks can generate multiple intermediate EMT states [10]. Additionally, mathematical models postulated stability of these states arising from intricate gene regulatory networks [10-12].

At the fundamental level, intermediate EMT states can be understood as either cell states *en route* to M-like states, or those stable states induced by weak (low-dose) EMT signals in the microenvironment. Recent single-cell transcriptomic studies showed that the time-dependent EMT programs contains intermediate states that delineate a continuum-like EMT spectrum [13-15]. However, it is unclear whether stable cell states in EMT program induced by multiple levels of signals support a continuum or a discrete EMT spectrum. While previous dose-dependent single-cell experiments with two EMT markers (E-cadherin for E, Vimentin for M) support the existence of intermediate EMT states [8, 10], much less is known about the transcriptomic profiles of the dose-dependent EMT spectrum.

In this work, we performed single-cell RNA-sequencing (scRNA-seq) using human mammary epithelial (MCF10A) cells treated with multiple concentrations of TGF-β. We found that the dose-dependent EMT program is a continuum containing multiple intermediate states that are stable after two-week treatment of TGF-β. We performed integrated analyses with our dataset and a recent time-dependent scRNA-seq dataset for the same cell line and EMT inducer [13] (**Figure 1A**). We found that the dose-dependent EMT spectrum has a stronger anti-correlation of E and M transcriptional programs than the time-dependent spectrum. While both spectrums show strong cell-to-cell variability and continuum-like patterns, the dose-dependent dataset has higher separability in terms of the groups of cells with neighboring labels (similar doses vs. similar time points). These differences enable higher sensitivity for the dose-dependent model to detect non-canonical EMT genes that are associated with the core EMT programs in terms of the expression pattern. Furthermore, the time-dependent dataset contains unique cell clusters at E-low region in the transcriptomic space, which correspond to *en route* cell states that do not appear at steady state. We found that signature genes in both dose- and time-enriched clusters are useful for prognostic predictions of cancer patients. Our analyses revealed key differences between dose- and time-dependent EMT programs in terms of the underlying dynamical processes, and showed the widespread existence of stable EMT continuum under multiple assumptions that may be relevant to physiological and pathological conditions.

**Figure 1.**
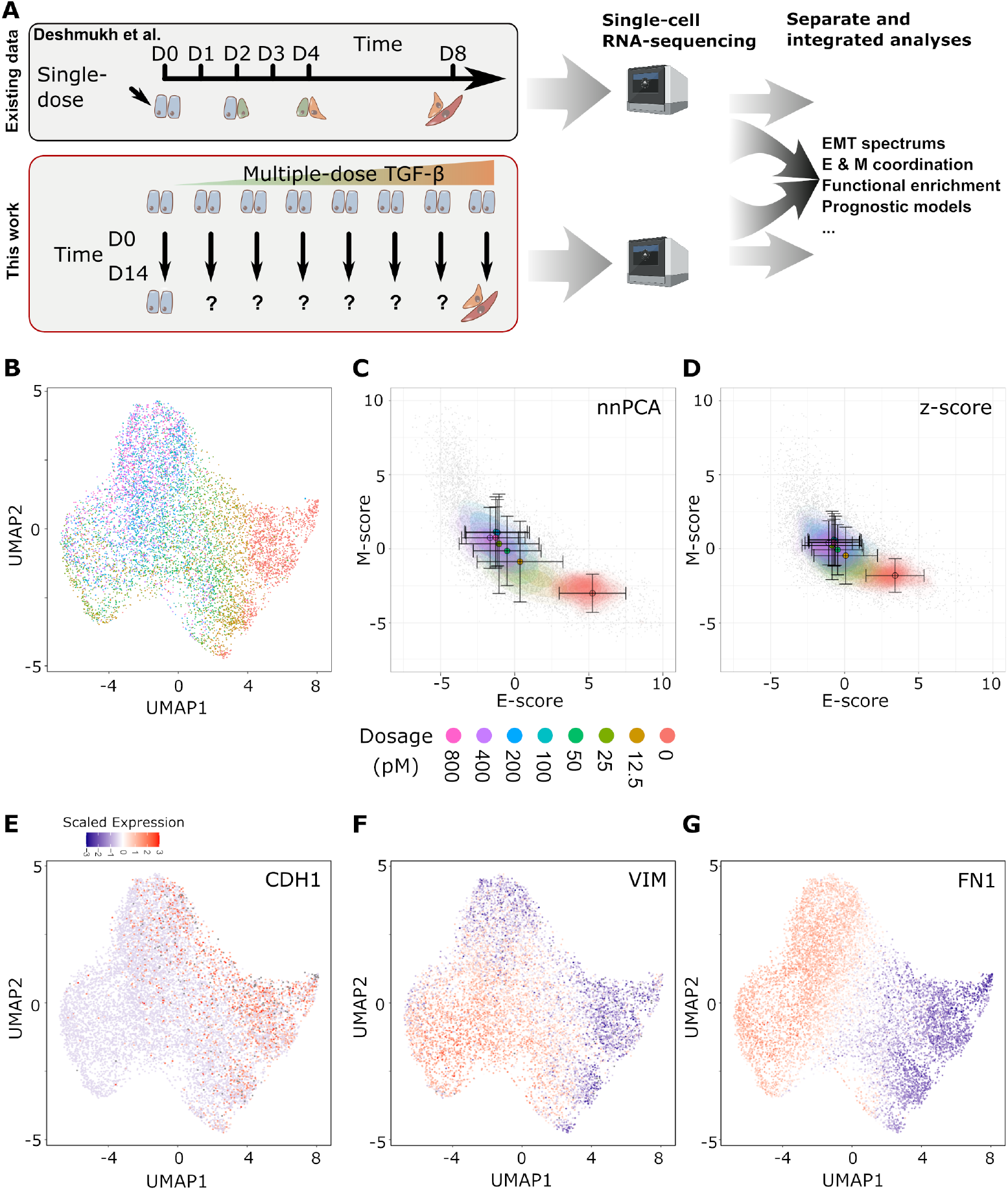
Analysis overview and progression of dose-dependent EMT at single-cell level. **(A)** A schematic of our analysis in this study. The analyses involve the existing time-course data (top) contain MCF10A cells at different time points following a common TGF-β treatment (about 200 pM; Deshmukh et al. [13]) as well as the dose-dependent data (bottom) containing MCF10A cells treated with different dosage levels of TGF-β after a fixed time period representing near-steady-state. Gene expression of cells from both experiments were measured using single-cell RNA-sequencing and were subsequently used individually and integrated for downstream analyses. **(B)** Projection of dose treatment single-cell expression data using UMAP. The color of individual points indicates the dose of TGF-β treatment from 0 pM (red) to 800 pM (pink). **(C-D)** Contour plots of gene set scores of E (x-axis) and M (y-axis) genes using nnPCA (C) and Z-score (D). Color indicates the dose of TGF-β as in (B). Circles indicate the mean E- and M-score of samples from each dose point and the associated error bars show the standard deviation (**E-G**) Overlay of the scaled expression of EMT marker genes CDH1 (an epithelial marker, E), VIM (a mesenchymal marker, F), and FN1 (a highly expressed mesenchymal gene, G). The color of individual points indicates the Z-score of expression of each gene from low (blue) to high (red).

## Results

### A single-cell transcriptomic landscape of dose-dependent EMT reveals a continuum-like spectrum

To characterize the transcriptomic spectrum with multiple levels of EMT signals, we performed dose-dependent induction of EMT with MCF10A cells, and analyzed cells at a near-steady-state time point (14 days after TGF-β treatment) using scRNA-seq (**Figure 1A, red box**). Transcriptomic profiles of 8876 cells with dosage annotation were identified after a standard filtering process and each condition of TGF-β concentration (dose) yielded more than 800 cells. We found that cells treated with various concentrations of TGF-β showed a continuous spectrum when visualized in the low-dimensional Uniform Manifold Approximation and Projection (UMAP) space (**Figure 1B**). To visualize the transcriptomic variability with interpretable, functional space, we used a recently developed projection method based on nonnegative principal component analysis (nnPCA) [16]. Previously identified epithelial-associated genes (E-genes) and mesenchymal-associated genes (M-genes), of which 203 E- and 136 M-genes were present in the processed dataset, were used to construct low-dimensional space with E-scores and M-scores (**Figure 1C**) [17]. We observed a progression of MCF10A cells from E-high-M-low state to E-low-M-high state with increasing concentrations of TGF-β. The effect of the progression was saturated with the TGF-β over 200 pM (**Figure 1C**). This progression was also observed with direct sums of the Z-scores of the E-genes and M-genes respectively with reduced resolution in terms of the separability of different conditions (**Figure 1D, Table 1**). Furthermore, we observed the continuous progression of key E-genes (e.g. CDH1) and M-genes (e.g. VIM and FIN1) expression (**Figure 1E-G**). The continuity of the transition and the saturation of the progression were similar to time-dependent EMT in MCF10A cells reported recently [13]. We also identified moderate divergence of mesenchymal-like cell states upon TGF-β treatment that appears to be driven by a difference in high (≥100pM) dose cells and predominantly lower dose cells in G1, which may have undergone cell cycle arrest [18] (**Figure S1**). Finally, we constructed Gaussian mixture models (GMMs) for the M-scores of all cells in the dose-dependent dataset, and we found that a non-binary, three-component model had the best fit to the data with highest scoring sub-optimal models favoring more, rather than fewer components (**Figure S2**). This dose-label-free approach suggests that the continuity of the spectrum was not merely due to the choice of TGF-β concentrations. Overall, our results show that the near-steady-state EMT program of MCF10A cells has a continuum-like spectrum that is independent of projection methods and sample labels.

**Table 1.**
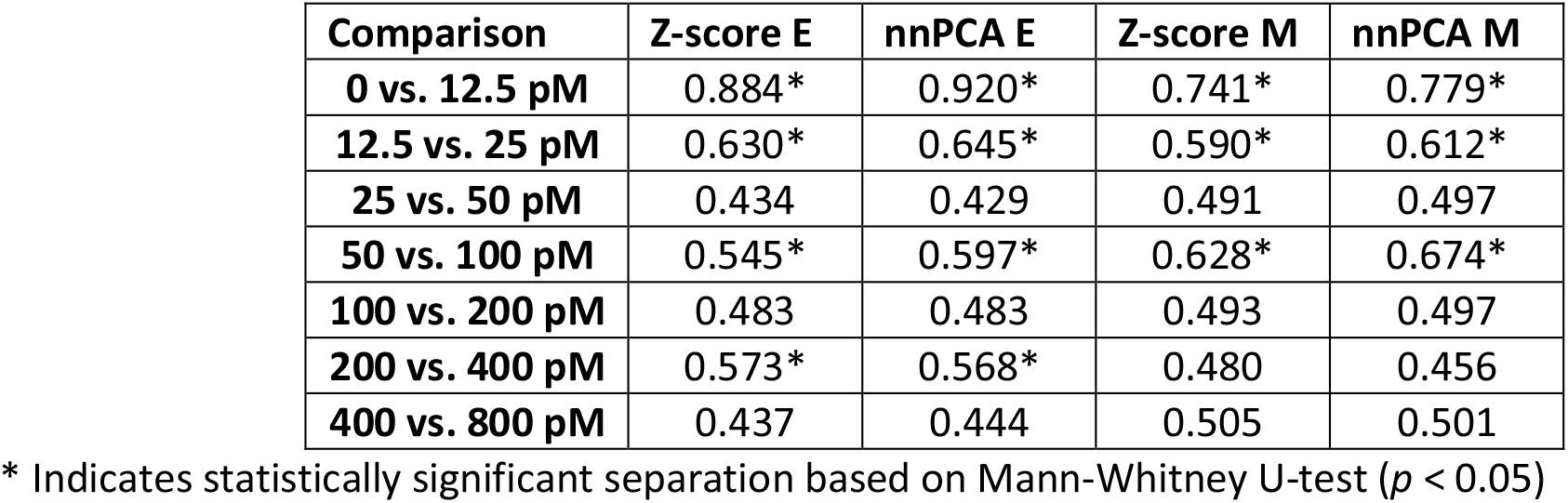
Common language effective size of separation of dosage groups based on EMT scores.

### Dose-dependent states show less variation and are more separable than time-dependent states in the E/M space

We next compared the dose-dependent EMT and time-dependent EMT at the single-cell resolution. We first integrated our dose-dependent dataset (abbreviated as dose dataset) with the previously published time-dependent dataset (abbreviated as time dataset) using the same approach as in Deshmukh et al. [13] (see **Methods)**. Because the time-dependent dataset contains two sets of experiment (short-time and long-time conditions), the integration involved three experiments. While comparison can be performed with unintegrated datasets, the integration is helpful as it reduces the effect of experimental batches not associated with the meaningful biological differences and increases the similarity between common cell states. As expected, we found that the integrated data contains cells from all experiments distributed in one region more continuously (**Figure 2A**) compared to the same analysis of unintegrated data (**Figure S3**). Furthermore, the untreated control in the dose-dependent dataset and the Time-0 control in the time-dependent dataset were located in similar regions in the expression space (**Figure 2A**, middle and right panels). These results suggest that the datasets can be compared in a reasonably uniform framework.

**Figure 2.**
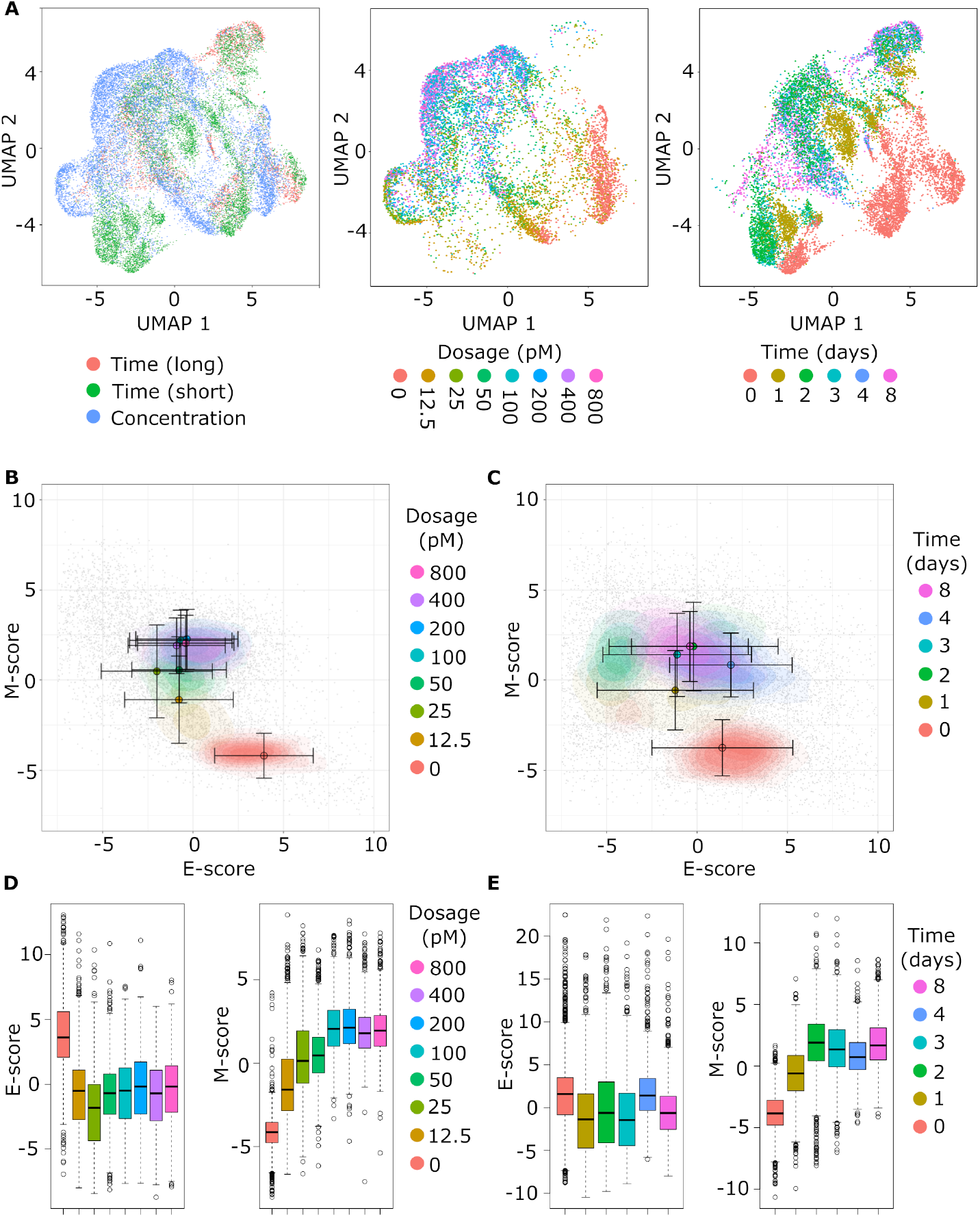
Continuity of integrated single cell dose and time data in low-dimensional projections. **(A)** Projection of integrated dose and time data using UMAP. Each panel uses the same underlying UMAP for samples, but the coloring of points has different meaning. Left panel: color indicates the origin of the sample from the Days 0, 4 and 8 of the time experiment (red), the Days 0, 1, 2 and 3 of the time experiment (green), or the dose experiment (blue). Middle panel: color indicates the treatment dose of samples from the dose experiment; time samples are masked. Right panel: color indicates the time of treatment for samples from the time experiment, dose samples are masked. **(B-C)** Contour plots of gene set scores of E (x-axis) and M (y-axis) genes using nnPCA for dose (B) and time (C) samples from integrate data. Color indicates the dose of TGF-β treatment from 0 pM (red) to 800 pM (pink) for dose data and time of treatment from 0 days (red) to 8 days (pink) for time data. Circles indicate the mean E- and M-score of samples from each dose point and the associated error bars show the standard deviation. **(D)** Boxplots show the distribution of E (left) and M (right) scores across different dose treatments from integrated data. Color indicates the dose of TGF-β as in (B). **(E)** Boxplots show the distribution of E (left) and M (right) scores across different time treatments from integrated data. Color indicates the time of treatment as in (C).

The data integration resulted in some alteration of the gene expression values and coverage (207 E and 141 M genes), but the progression of EMT in the dose-dependent manner was preserved, especially in the mesenchymal axis (**Figure 2B and D**). Notably, the time-dependent data points were distributed more broadly compared to the dose-dependent data (**Figure 2A-C**). Nonetheless, the continuous E-to-M progression was observed (**Figure 2C and E**). Note that the broader distribution of cells in the time-dependent dataset was also observed with unintegrated data (**Figure S4**). Dosage data maintained a similar pattern of separability as the unintegrated data with superior differences between conditions up to the putative saturation point between 100 and 200 pm. Time samples, however, show greater separability between the earliest (0 vs 1 day) and latest (4 vs 8 days) time points, with the separation of interior points varying between E and M scores and failing for 3 vs 4 days, which is the breakpoint between the two time data batches (**Table 2**).

**Table 2.** Common language effect size of separation of time and dosage groups based on EMT scores of integrated data.

| Comparison | nnPCA E | nnPCA M |
| --- | --- | --- |
| **Time** |  |  |
| **0 vs. 1 days** | 0.685* | 0.883* |
| **1 vs. 2 days** | 0.439 | 0.781* |
| **2 vs. 3 days** | 0.554* | 0.439 |
| **3 vs. 4 days** | 0.281 | 0.416 |
| **4 vs. 8 days** | 0.701* | 0.650* |
| **Dosage** |  |  |
| **0 vs. 12.5 pM** | 0.890* | 0.905* |
| **12.5 vs. 25 pM** | 0.618* | 0.688* |
| **25 vs. 50 pM** | 0.377 | 0.532* |
| **50 vs. 100 pM** | 0.487 | 0.753* |
| **100 vs. 200 pM** | 0.469 | 0.517 |
| **200 vs. 400 pM** | 0.554* | 0.434 |
| **400 vs. 800 pM** | 0.448 | 0.523* |

\* Indicates statistically significant separation based on Mann-Whitney U-test ( $p < 0.05$ )

In addition to the difference in the width of the distribution, the dose-dependent dataset had stronger anti-correlation between E and M scores (*R* = −0.531) than the time dataset (*R* = −0.288) **(Figure 2B and C**). We hypothesized that the stronger coordination between E and M transcriptional programs can facilitate the discovery of EMT-associated genes within our integrated dataset that were not classically considered EMT genes in previous studies [17, 19]. Indeed, with the same threshold of Pearson correlation coefficient (0.25), the dose-dependent dataset revealed greater numbers of genes that had significant association (correlation with M-scores and anti-correlation with E-scores or vice-versa) with the overall expression of E and M genes (191) compared to the time-dependent dataset (42) (**Figure 3A and B**). The genes that showed significant association with E-program in the dose-dependent dataset but not in the time-dependent dataset were enriched with functions such as keratinization, keratinocyte differentiation and epidermis development, while M-correlated genes included those associated with integrin and chemokine binding (**Figure 3C**). Notably, TGF-β has been shown to be involved in keratinocyte growth arrest [20], which is consistent with keratinocyte differentiation being associated with the E-program. We then focused on specific genes there were specifically correlated in with EMT progression in dose data, but not time data. FARP1 was among the genes most highly correlated M-scores scores (*R* = 0.57) in dose data, but not in time data (*R* = 0.17). FARP1 was recently shown to be important for cancer cell motility and associated with poor prognosis [21]. Similarly, ESM1 was highly correlated with M-scores in dose (*R* = 0.50) but no time (*R* = 0.10), and it contributes to the metastasis in colorectal cancer via NK-ƙB activation [22]. Note that these analyses were performed in the absence of dose and time treatment labels, so the correlations were primarily driven by the intrinsic EMT continuum. Our results suggest that compared to the time-dependent data, the dose-dependent scRNA-seq data may provide higher sensitivity to detect non-classical EMT genes that are coordinated by the core EMT module.

**Figure 3.**
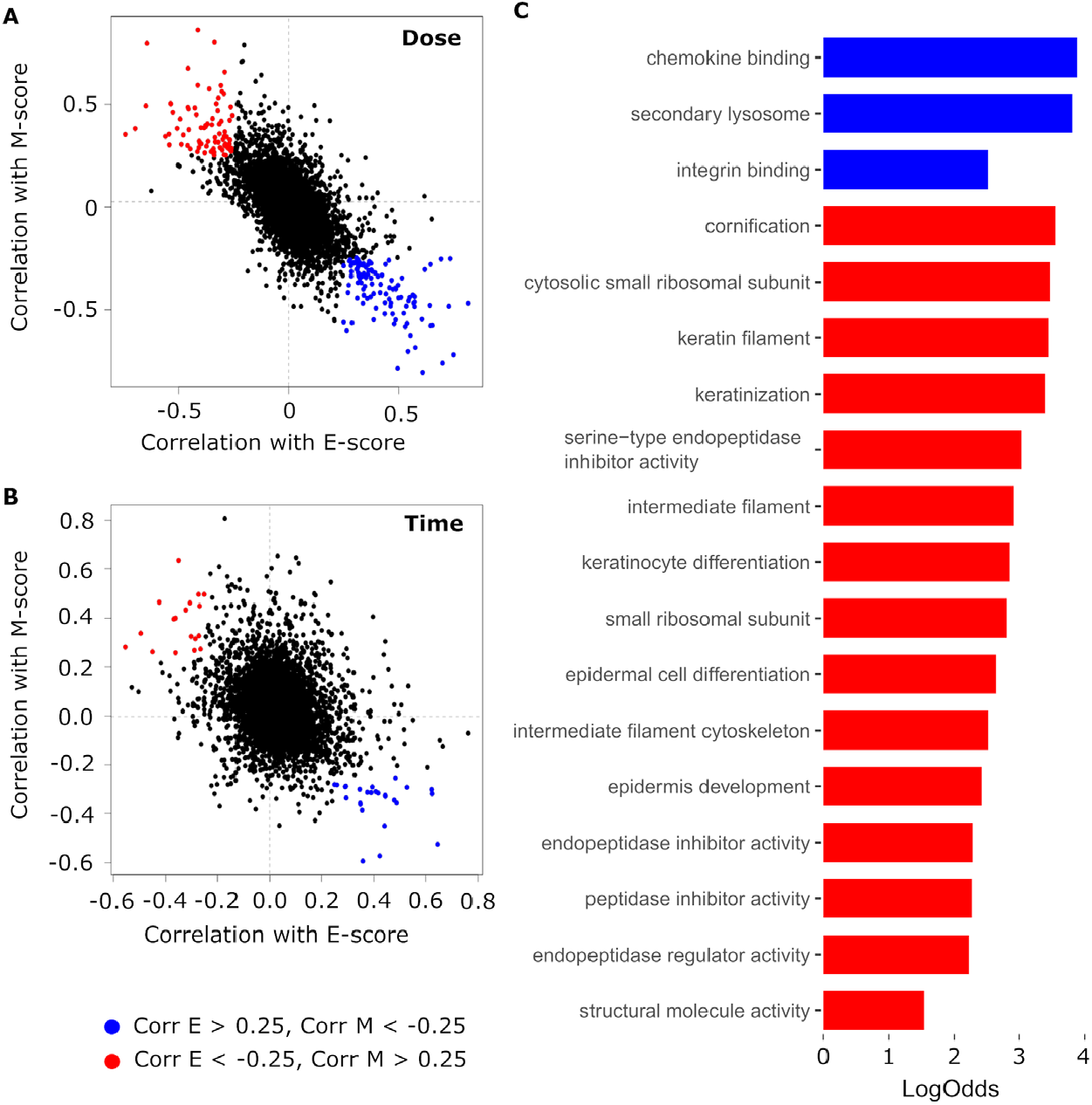
Correlation of non-classical EMT genes with E and M scores in integrated data. **(A-B)** Correlation (*R*) of non-EMT gene expression with sample E-scores (x-axis) and M-scores (y-axis) in dose (A) and time (B) samples from the integrated dataset. Each point represents one of the 15000 most variable genes from the integrated single cell dataset, excluding those genes used to define the E- and M-scores. Blue points indicate genes which are anti-correlated with EMT progression (*R* > 0.25 for E-scores, < -0,25 for M-scores), while red points indicate genes which are correlated with EMT progression (*R* < -0.25 for E-scores, > 0.25 for M-scores). **(C)** Bar chart showing the log odds of the top 5 biological process, molecular function, and cellular component GO terms enriched in EMT correlated (red) and anti-correlated (blue) genes in dose data. GO terms were selected based on adjusted *p*-value and if there were less than 5 GO terms in each category with an adjusted *p*-value < 0.05, only those terms with an adjusted *p*-value < 0.05 were reported.

### *En route* cell clusters unique to time-dependent EMT program

Based on the distinct distributions of cells between time- and dose-dependent EMT programs (**Figure 4A**), we wondered whether our comparative study can reveal expression regions containing cells that are *en route* to the M state rather than (partially) stabilized at intermediate EMT attractors. To simplify the representation of the expression profiles in the EMT spectrum, we focused on a 4 ×4 grid in the E- and M-score space. We found that a region with low scores of one (E or M) program, and low-to-medium scores of the other program (M or E) is enriched with cells in the time-dependent EMT program (**Figure 4B, lower left**). This result reveals a transient EMT path that may primarily involve the relatively low expression of both E and M genes. Nonetheless, the enrichment of time data points in the regions of high E-gene activity and M-gene activity suggested a possible alternative transient EMT path (**Figure 4B, right**) [23, 24]. We then identified significantly differentially expressed genes across all sixteen segments of the 4 ×4 grid and focused first on known EMT marker genes. Interestingly, while the transient path crossing the E-low-M-low region is generally consistent with the profiles of the E marker CDH1 and EPCAM as well as M markers VIM and FN1 (**Figure 4C-F**), a distinct sequence of M gene activations was observed in the hypothetical E-to-M path: for example, VIM was activated before FN1 in this path (**Figure 4E and F**). This observation is consistent with our earlier transcriptomic data showing significant diversity of M-genes in response to EMT signals [19]. We also found correspondence between the time versus dose enrichment and graph-based clusters of the full data (**Figure S5)**. Two clusters containing predominantly (>75%) time samples correspond to the transient low expression path while another cluster overlaps with time-enriched segment with high E but moderate M expression (**Figure S5B**). Similarly, two clusters containing predominantly dose samples further show that dose is enriched in the most extreme high E, low M and low E, high M segments, while a third cluster reflecting the general enrichment of dose samples in high M segments towards the middle of E-and M-score space (**Figure S5C**). We then divided the 4 ×4 grid into time and dose enriched regions based on log-transformed odds ratio (LogOdds), and looked for Gene Enrichment (GO) terms among significantly up-regulated genes. Time-dominant segments had LogOdds greater than 0.4, dose-dominant segments had LogOdds less than -0.4, and segments without and differentially expressed genes were not considered for the subsequent GO analysis (**Figure 4G**). Overall, we identified 1378 enriched biological process and molecular function terms enriched in at least one segment. 102 of the terms were predominantly found in either the time enriched or dose enriched region (≥ 50% of segments in the regions and twice as many segments as the opposing region). Among the terms with the greatest differences in enriched segments between regions include ‘extracellular matrix structural constituent’ and ‘endodermal development’ (which favor dose) as well as ‘neuron death’ and ‘cellular detoxification’ (which favor time, **Figure 4G**)

**Figure 4.**
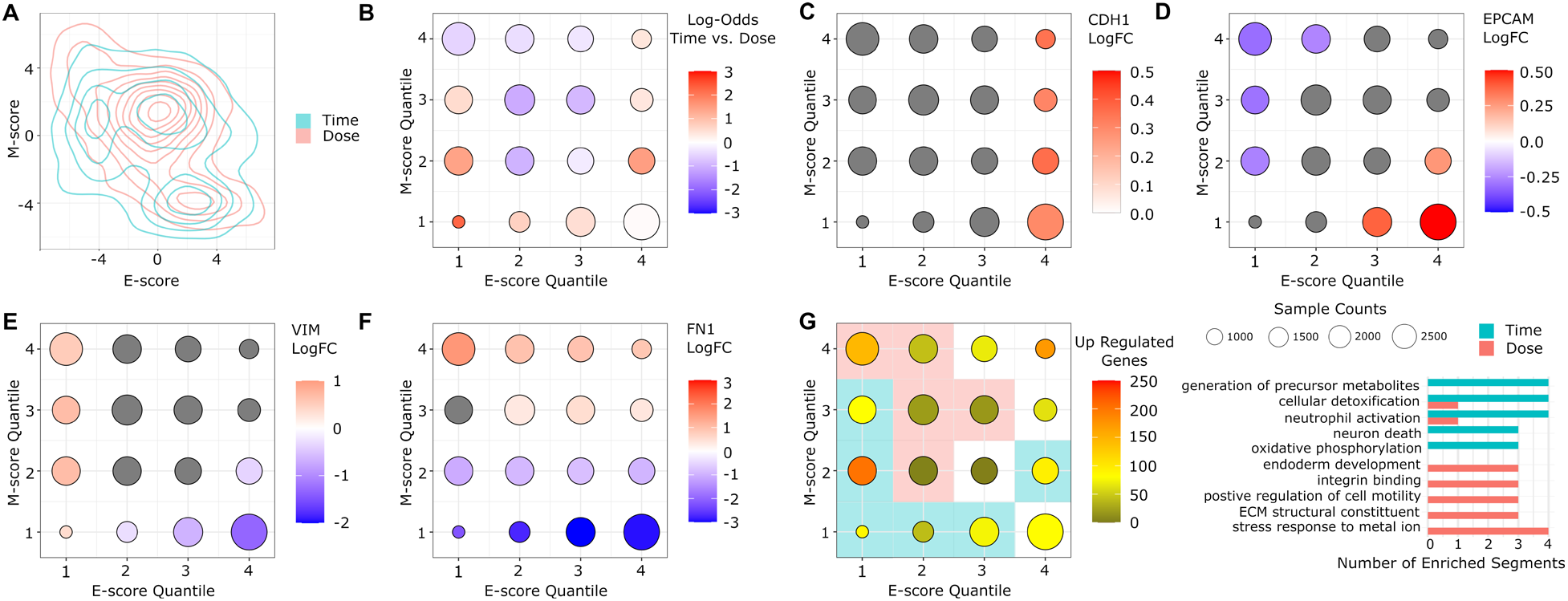
Enrichment of dose and time samples in E- and M-score space. **(A)** A contour map showing the distributions of time (blue) and dose (red) samples in E- and M-score space. **(B)** Bubble chart showing the enrichment of time samples in different segments of E- and M-score space. Each point represents a segment of E- and M-score space defined by a particular quartile of E-score (x-axis) and M-score (y-axis). The size of the point corresponds to the total number samples in the segment and the color of the point represent the log-odds of time sample enrichment from low (blue) to high (red) with a log-odds of 0 (white) indicating balanced representation. (**C-F**) Bubble charts showing the differential expression of key EMT genes, CDH1 (C), EPCAM (D) VIM (E), and FN1 (F) in different segments of E- and M-score space. Point positions and size are defined as in (B). The color of the point represents the log fold change of expression of the gene from low (blue) to high (red) with a log fold change of 0 (white) indicating no-change relative to other samples. Gray circles indicate below-threshold (± 0.25 log fold change) differences. (**G)** Left: A bubble chart showing the number of significantly up-regulated genes in each segment of E- and M-score space. Point positions and size are defined as in (B). The color of the point represents the number of upregulated genes from 0 (black) to > 200 (red). The colored shading behind the points represents whether the segment is enriched from time samples (log odds > 0.4, blue) or dose samples (log-odds < - 0.4, red), ignoring segments without any upregulated genes. Right: Bar chart showing 5 GO terms enriched across time and dose segments. GO term were chosen based on the difference in number of enriched segments between time and dose and distinction function. Significant enrichment was assessed based on adjusted *p*-value (*p* < 0.05).

### Signature genes from both dose- and time-dependent datasets contribute to better prognostic models

To examine the roles of the signature genes at various locations of the EMT spectrum in prognosis, we again focused on the sixteen segments of the 4 ×4 grid of the EMT space containing both time and dose data. We used 27 diverse cancer datasets with tumor RNA-seq and patient data from The Cancer Genome Atlas (TCGA). We used the signature genes of each segment to construct a Lasso penalized Cox model for predicting the survival outcomes of cancer patients for each of the 27 cancer types. Interestingly, signature genes in both the E-low-M-high segment corresponding to the extreme M-state, and some intermediate EMT segments had relatively high average prognostic performance measured in C-index (**Figure 5A, red**). Notably, the high-performing intermediate EMT segments had either low E-scores or high M-scores. We scaled performance index across cancer types to account for dataset level variance in model construction, (**Figure 5B**). We found that, in general, E-low intermediate states (green) and M-high intermediates (light blue) both outperformed other (blue) segments with similar average performance to the extreme M state (yellow), although there was a large degree of variance across cancers and on the high end of the performance spectrum. The superior prognostic performance of the extreme M-state and certain intermediate states was also observed in the recent study based on time-dependent EMT alone [13], and our analysis revealed the characteristics of these high-performing groups in the 2-dimensional EMT spectrum. Unsurprisingly, the performance of the E-low segments was primarily driven by the time data (**Figure 5C**), whereas both time and dosage data contributed to the performance M-high segments (**Figure 5D**). The low performance of the E-medium-M-medium segments was at least partially due to the low numbers of signature genes in these groups (**Figure 5A and C, middle**). Nonetheless, the condition (time or dose) plays a significant role in the two E-high segments which only perform well in dose data (**Figure 5A and D, right**).

**Figure 5.**
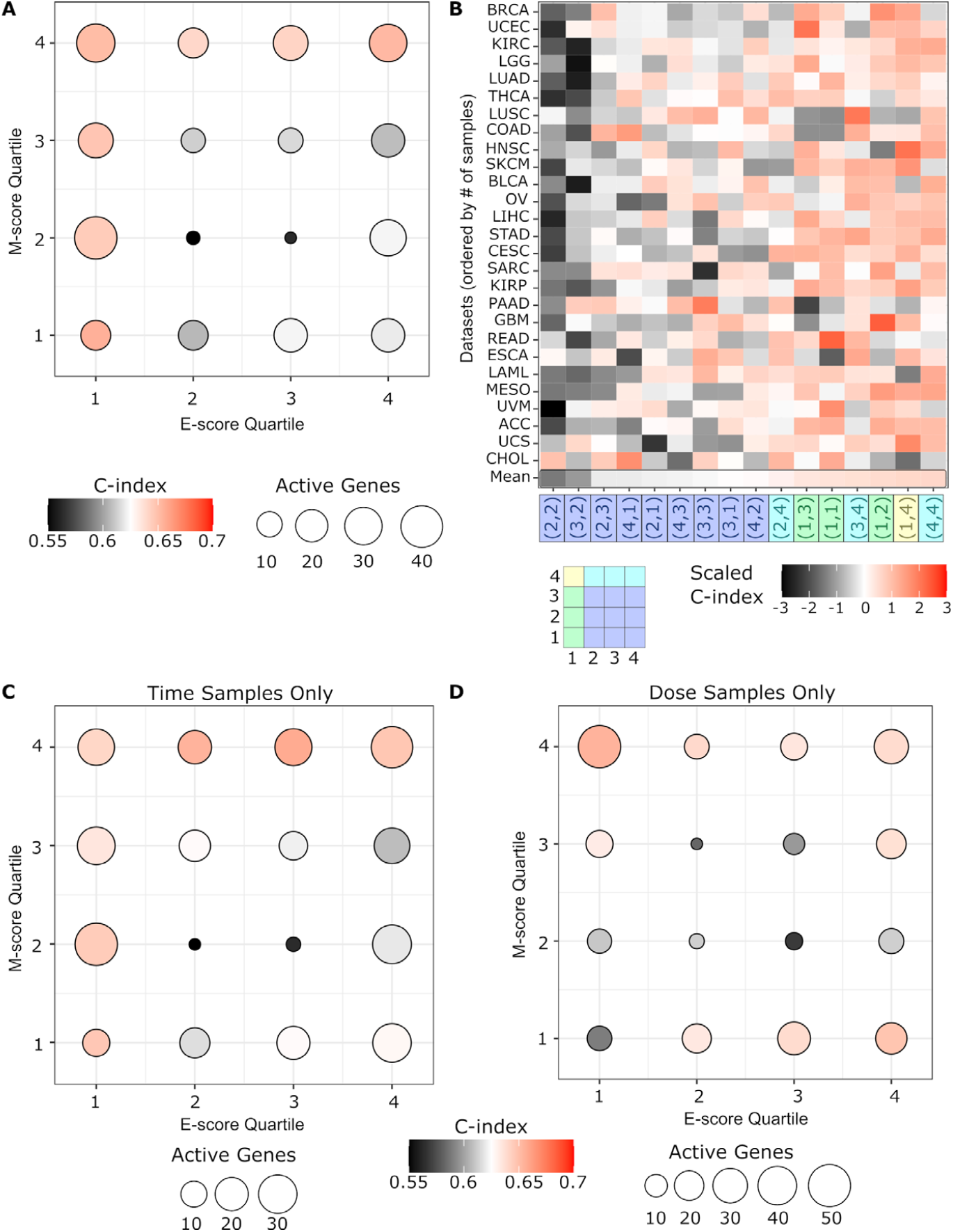
Performance of prognostic models. **(A)** Bubble chart shows the average performance of Cox hazard models built using the up-regulated genes from different segments of the E- and M-score space. Each point represents a segment defined by a particular quartile of E-score (x-axis) and M-score (y-axis). The size of the point corresponds to the average number predictor genes across all models built using the up-regulated genes from that segment. Color represents the C-index averaged across all models of cancer datasets, where C-index is a measure of the correspondence between the models predicted risk and survivorship, similar to the Area Under the Receiver Operating Curve (AUC-ROC). Red represents higher C-index (better performance). Black represents lower C-index (poorer) performance. White represents the approximate median of scores (0.635). **(B)** Heatmap of individual model performance across all combinations of segment-gene-derived models (x-axis) and cancer datasets (y-axis). The color of individual cells represents the C-index of the model scaled against all models of that cancer datasets to account for data-set-level differences in model performance. Segments are ordered across the x-axis by the increased average scaled C-index from left to right, while cancer datasets are ordered by increasing number of samples from bottom to top. The shading of segment coordinates along the x-axis distinguishes between the extreme M state (yellow), E-low intermediates (green), M-high intermediates (light blue), and other segments (dark blue). **(C-D)** Bubble charts show the average performance Cox hazard models built using the up-regulated genes identified using only time (C) or dose (D) samples from different segments of E- and M-score space. Point position, size, and color as defined as in (A).

## Discussion

Time dependent EMT processes have been extensively studied with single-cell transcriptomics in recent years [3, 13, 14]. While these studies provided substantial information about EMT progression in multiple contexts, the connection between the intermediate cell states observed in these experiments and cell attractors was elusive. Using near-steady-state single-cell transcriptomic profiling, we showed that the EMT spectrum can be described as a stable continuum under multiple levels of EMT signals. This information complements earlier studies with single-dose time course data, and it provides stronger evidence supporting the existence of multiple intermediate EMT states that widely exist in tumors and metastatic cells [5]. Furthermore, by comparing the time- and dose-dependent single-cell data, we identified groups of cells that are exclusively *en route* to M state, which shows the possibility that some cells can transiently deactivate the epithelial program before activating the mesenchymal program. Nonetheless, it is likely that many other *en route* cells are also close to the stable intermediate EMT states, and these states have intermediate levels of both E and M genes. Furthermore, given the heterogeneous microenvironment of cells under physiological conditions, the cells states may be determined by both the multi-level EMT signals and the time-dependent stages of the EMT process, so both dose dependent and time dependent *in vitro* data can contribute to the understanding of the EMT program *in vivo*.

Previous mathematical models have provided mechanistic insights into the gene network structures supporting multiple intermediate EMT states [10-12, 25-28]. While interconnected positive feedback loops involving a few genes can govern discrete intermediate states, the continuum-like, wide distributions of cells in the EMT spectrum even under the near-steady-state condition suggest that the discrete cells states may only partially explain the stable phenotypical heterogeneity. At least two mechanisms may explain the gap between the existing theories the observed continuum: realistic gene regulatory networks may contain much more positive feedback loops than those described by existing models [29], and these loops can support many intermediate states; dynamical and reversible cell-state transitions at stationary phase, such as those driven by transcriptional noise and slow-timescale oscillations [30], can give rise to cells that are far away from point attractors in the gene expression space. Nonetheless, existing models have provided a strong theoretical foundation for stable intermediate EMT states, and future development of the theories and quantitative experiments will be important to the understanding of EMT continuum.

EMT involves dramatic phenotypic changes of cells. It is therefore expected that transcriptome-wide alteration is induced during the multi-stage transition. This suggests that classically defined core EMT genes may not be sufficient to provide a holistic view of the EMT program. We showed that the dose-dependent single-cell transcriptomes can be useful to identify genes that show expression patterns highly correlated with the core EMT genes at the single-cell level. The dose-dependent scRNA-seq experiment can therefore provide crucial links of the core EMT networks to the rest of the transcriptomes, and some of these connections may not be revealed by time-course scRNA-seq experiments due to the large numbers of cells that transiently activate parts of the transcriptional program. Nonetheless, the two types of scRNA-seq experiments contain complementary information, and we suggest that both can be used in future studies to reveal cellular programming that determines cell-to-cell variabilities in cell populations.

## Methods

### Cell culture

MCF10A cells were obtained from ATCC and grown in DMEM/F12(1:1) medium with 5% horse serum, epidermal growth factor (10 ng/mL), cholera toxin (100 ng/mL), and insulin (0.023 IU/mL). For TGF-β treatment, cells were incubated with indicated concentrations (**Figure 1B**) of human TGF-β1 protein (R&D systems) in the complete culture medium. The culture medium was replaced daily, and cells were passaged right before reaching full confluency.

### Single-cell RNA-sequencing

MCF10A cells were first labelled with Perturb-seq vectors without sgRNA expression using guide barcodes (GBCs) that were originally used to identify sgRNAs [31]. Barcoded MCF10A cells were then treated with different dosages of TGF-β for 14 days and single cells were prepared and mixed at a concentration of approximately 1,000 cell/μL. Transcriptome library generation was performed following the Chromium Single Cell 3' Reagents Kits v2 (following the CG00052 Rev B. user guide) where we target 10,000 cells per sample for capture. GBC library was generated from a fraction (5ng) of amplified whole transcriptome by dial-out PCR method according to a previous publication [31]. Both libraries were mixed at 9:1 ratio and sequenced by paired-end sequencing (26bp Read 1 and 98bp Read 2) with a single sample index (8bp) on the Illumina HiSeq 2500. Generated FASTQ files were aligned utilizing 10×Genomics Cell Ranger 2.1.0. Each library was aligned to an indexed hg38 genome using Cell Ranger Count. The cell barcode (CBC)-GBC table was generated from the GBC library and used to identify the treatment groups.

### Data processing and integration

Sequencing data for time-course single-cell data from Deshmukh et al. [13] was obtained from National Center for Biotechnology Information Sequence Read Archive (BioProject ID: PRJNA698642) and mapped using to the same human genome assembly as our dose (CRXh38.84) data using Cell Ranger [32]. Aligned sequences we processed using the Seurat package (version 4.0.2) in R [33]. Gene names between experiments were correlated using the HGNChelper package [34], using the suggested gene symbol for each gene except when it would create a duplicate reference. Genes were filtered from individual runs if they did not appear in three or more cells. We then filtered each dataset for cells with fewer than 500 features or more than three median absolute deviations beyond the median number of features of the sample set (i.e., the long time, short time, and dose datasets). We additionally eliminated any sample with a fraction of mitochondrial reads that was greater than 0.2. We then integrated all time and dose data following the procedure used in Deshmukh et al. to best preserve the relationship between time samples observed in the study [13]. Briefly, the integration involves identifying the most variable genes in each dataset, defining ‘anchor’ samples between datasets using canonical correlation analysis, correcting the expression of related anchor samples and finally propagating this correction to other samples based on the similarity to the anchors. We normalized and calculated cell cycles scores for each dataset independently prior to integration and used the 15000 most variable genes to identify anchors. We applied the same top 15000 variable genes filter to unintegrated dose data for calculating UMAP, nnPCA scores and clustering. Finally, each dataset was scaled across genes and the expression was corrected for cell-cycle phases.

### Projection of single cell data in reduced dimensional space

Projection of single cell data was done using the scaled expression values for both unintegrated and integrated data. For this process we employed three main approaches, Uniform Manifold Approximation and Projection (UMAP), average Z-scores, and non-negative Principal Component Analysis (nnPCA). For UMAP, we performed principal component analysis (PCA). We then used the first 15 principal components to construction the map using the RunUMAP function from Seurat with defaults parameters beyond specifying the PCA input and size. UMAP was primarily used to assess the continuity of both unintegrated and integrated datasets. Average Z-score was determined using the gsva package [35]. nnPCA-based scores were computed by performing non-negative principal component analysis on subsets of the scaled dataset defined using epithelial (E) and mesenchymal (M) genes identified by Tan et al. [17]. We used the nsprcomp function from the R package of the same name with the option nneg=TRUE and ncomp=5 to identify the top five components for each subset [36]. While the first five components we calculated, only the first was used as the function greedily optimizes the variance explained by each component in order. Further details about our nnPCA scoring approach can be found in Panchy et al. [16].

### Gaussian mixture models

Gaussian mixture models of E and M scores were performed using the mclust package [37] and figures were generated using the plotMix function from MineICA [38]. For each score, we used the unequal variance model (V) with the optimal Bayesian information criterion (BIC) as our optimal model.

### Enrichment analysis

To examine the distribution of time and dose samples, we divided E and M-score space into a 4 ×4 grid based on the 25^th^, 50^th^ and 75^th^ percentile of E and M-scores. Our dose dataset contains 2402 unlabeled dose samples (within a reasonable range [31]), which show roughly even distribution across unintegrated dose data in both UMAP and E- and M-score space (**Figure S6**), suggesting that they are merely missing annotation rather than contamination. Nonetheless, to obtain results comparable to progression plots in **Figure 2**, enrichment only considered labeled time dosage samples. Odds of the enrichment of time and dose samples in each segment of the 4 ×4 grid of E and M-score space were calculated using the Fisher’s exact test implemented in R (fisher.test). Differentially expressed marker genes for each segment were identified using the FindAllMarkers function from Seurat. As recommended, for FindAllMarkers we used unintegrated, normalized counts and introduced non-independence between samples. For GO enrichment in each segment, we selected all genes in each segment with a fold change in expression > the 75^th^ percentile of all marker genes across all segments and an adjusted *p*-value < 0.05. GO enrichment of identified genes set was performed using the clusterProfiler package in R [39], using a background of all expressed genes in the unintegrated data, and significance was assessed using the Benjamini-Hochberg adjusted *p*-value.

### Prognostic models

TCGA bulk RNA-seq data and associated meta-data were obtained from TCGAbiolinks [40]. Of the 33 available datasets (cancer types), we selected those with at least ten samples in both the survivor and non-survivor groups, which gave rise to the 27 samples listed in **Figure 5B**. For each dataset, FPKM expression values from all non-normal tissue samples were extracted. For features, we used the set of up-regulated marker genes (a fold change in expression > the 75^th^ percentile of all marker genes across all segments and an adjusted *p*-value < 0.05) in each segment. Each combination of cancer expression data and gene set was then used to construct Cox proportional hazard models using glmnet implemented in R. In brief, a glmnet model employed Lasso regularization (alpha = 1) and performed 10-fold cross validation to optimize the lambda parameter which influences the strength of regularization. In theory, the stronger Lasso regularization should reduce the number of active predictors (genes) in the model, but we do not actively seek the minimized predictors. Instead, we report the model which maximizes the average cross-fold C-index, which is the proportion of concordant pairs to total pairs in the dataset, i.e. the proportion of all possible samples pairs where increased model hazard corresponds to reduced survival. This measure is roughly analogous to AUC-ROC in a censored data context such as survival.

### Clustering from shared-nearest-neighbor graphs

To confirm that the associations we found in E- and M-score space reflect the full dataset, we examined sample associations across the 15000 most variable expressed genes in the integrated and dose only dataset using the clustering based on shared-nearest-neighbor graphs via Seurat. After scaling and correcting single cell data, principal component analysis was done using RunPCA with 15 components (npcs=15) and then passed to FindNeighbors with options reduction=‘pca’, dims=1:15. Finally, FindCluster was run with resolution = 0.5 and default parameters and seeding otherwise.

## Data and code availability

All code and sequencing data are available at GitHub repository: https://github.com/panchyni/Time_and_Dose

GEO: GSE200941

## Ethics declaration

All experimental work was performed in compliance with protocols approved by RIKEN Center for Integrative Medical Sciences. No human, animal subject, tissue, gamete, or stem cell was involved in this study.

## Author contributions

Designed research: KW and TH. Performed experiments: KW and MT. Analyzed data: NP, KW, AW and TH. Wrote manuscript: NP, KW and TH. All authors read and approved the manuscript.

## Supplemental Figures

**Figure S1.**
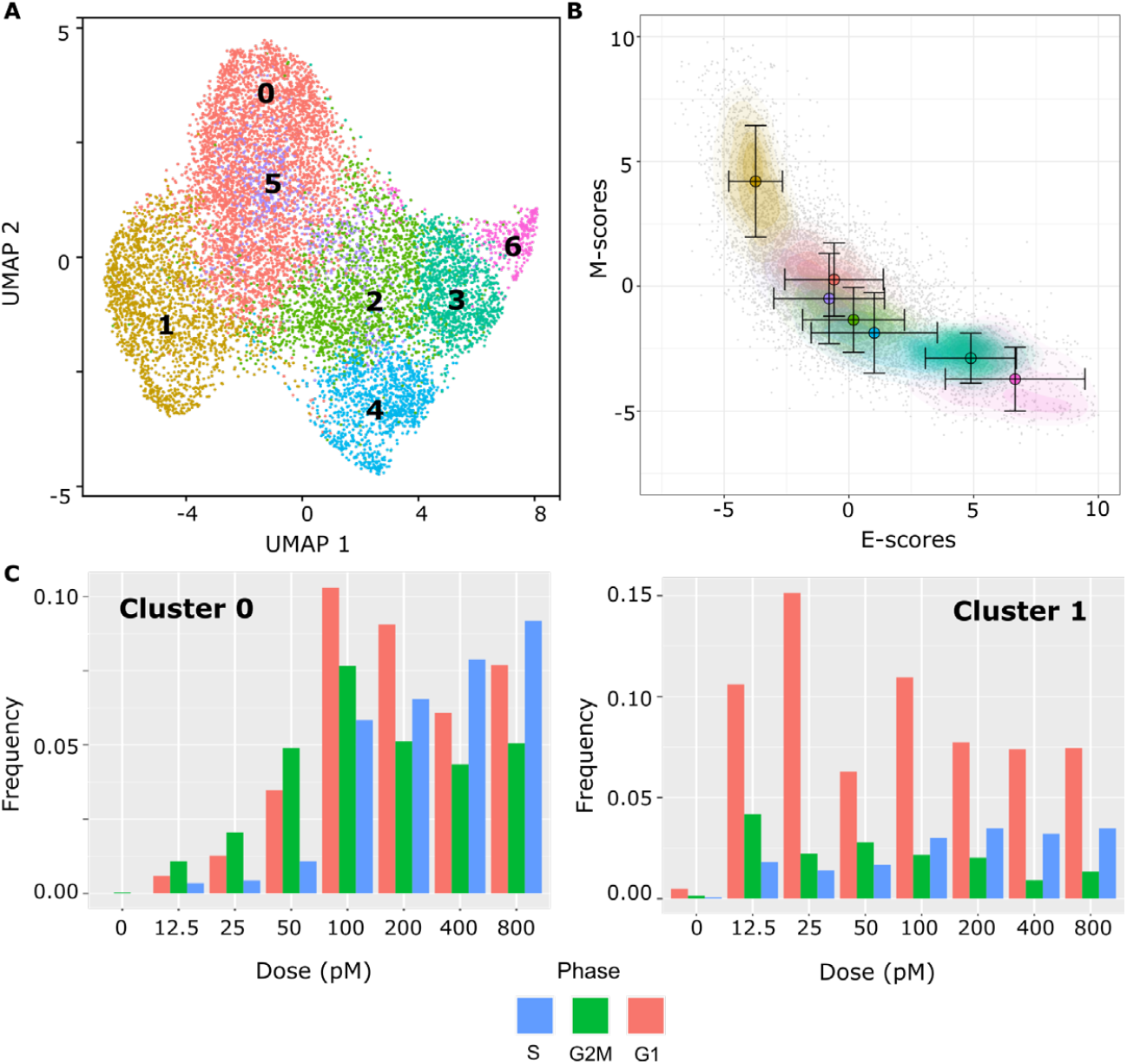
Clustering based on shared-nearest-neighbor graphs in dose UMAP and E- and M-score space. **(A)** Projection of dose treatment single cell expression data using UMAP. The color of individual points indicates the clusters based on shared-nearest-neighbor graphs to which the sample was assigned **(B)** Contour plots of gene set scores of E (x-axis) and M (y-axis) genes using nnPCA for dose data. Color indicates cluster of samples in UMAP space, with individual colors corresponding directly to clusters in (A). Circles indicate the mean E- and M-score of samples from each dose point and the associated error bars show the standard deviation. **(C)** Bar charts show the frequency (y-axis) of different treatment dosages (x-axis) in Cluster 0(left) and Cluster 1 (right). The frequency of each dosage is broken down by the inferred cell-cycle state: S (blue), G2/M (green) and G1 (red).

**Figure S2.**
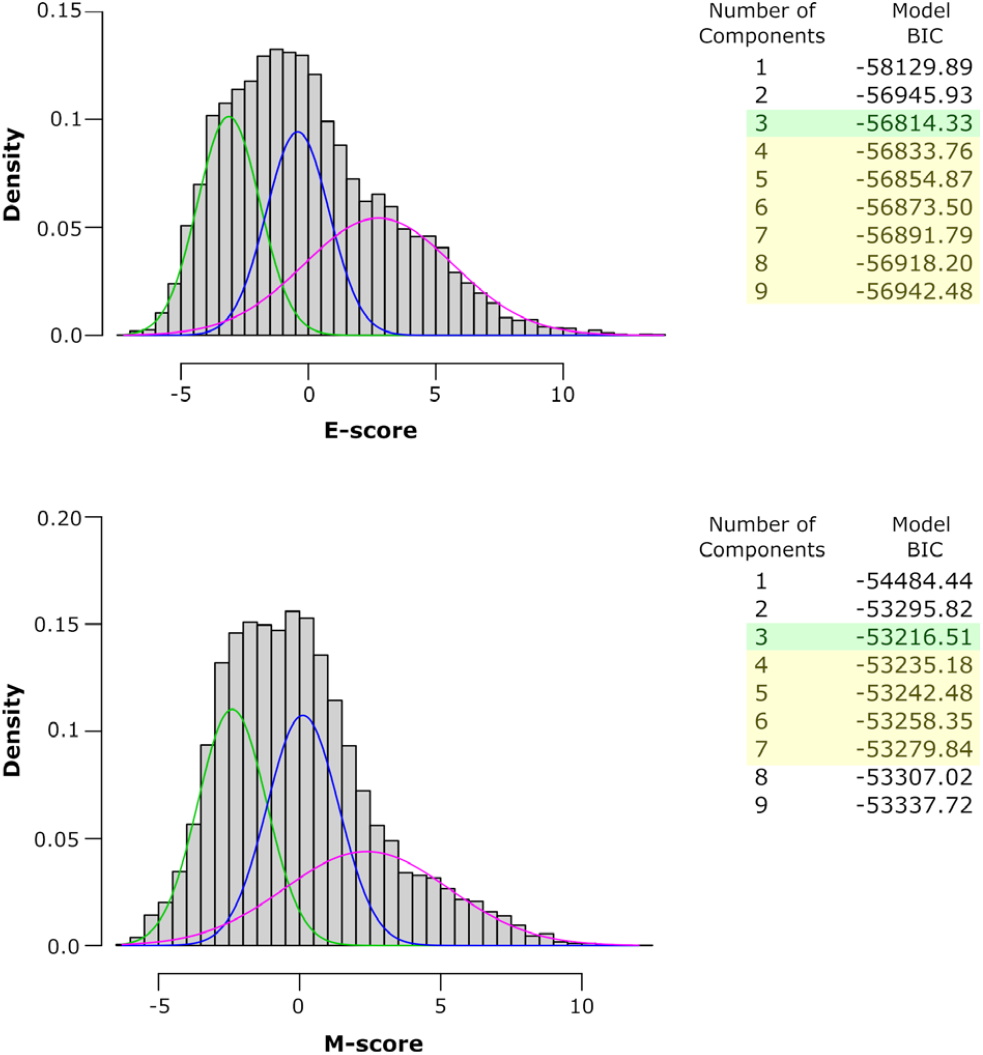
Gaussian mixture model of E- and M-score of dose data. Gaussian mixture model (GMM) of E-score (top) and M-scores (bottom) of dose treatment single cell data. Each panel shows a histogram of scores with the score values along the x-axis and density along the y-axis. The colored curves overlain on the histogram show the number, position, and spreads of the optimum GMM for each score. To the right of each histogram is a table of BIC scores for GMM models with a different number of components with the optimal model highlighted in green and sub-optimal models with better fits than a binary model highlighted in yellow. Note that McClust calculates the inverse of the normal BIC score, so the maximum score indicates the best fit.

**Figure S3.**
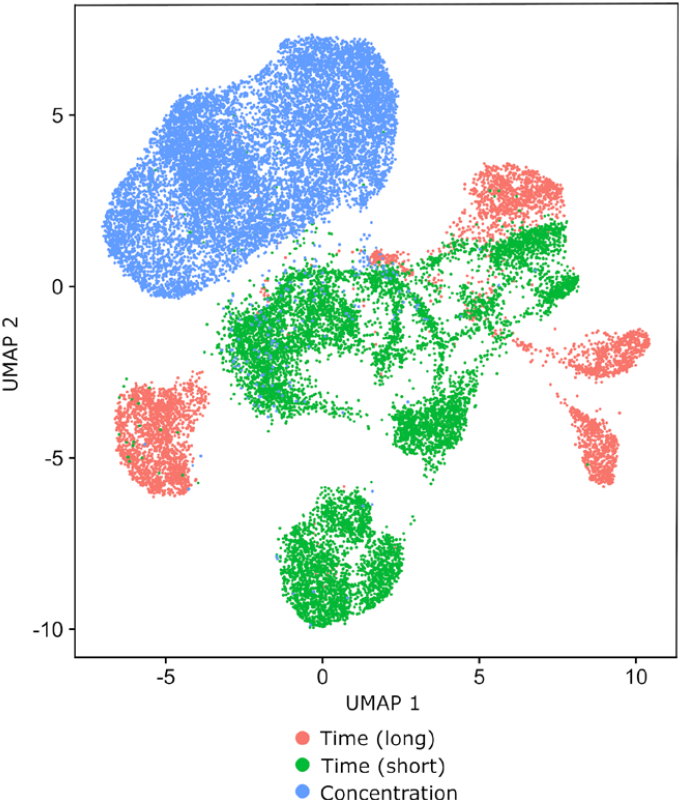
Positioning of samples from different experiments in unintegrated UMAP Space. Projection of unintegrated dose and time data using UMAP. Color indicates the origin of the sample from the Days 0, 4 and 8 of the time experiment (red), the Days 0, 1, 2 and 3 of the time experiment (green), or the dose experiment (blue).

**Figure S4.**
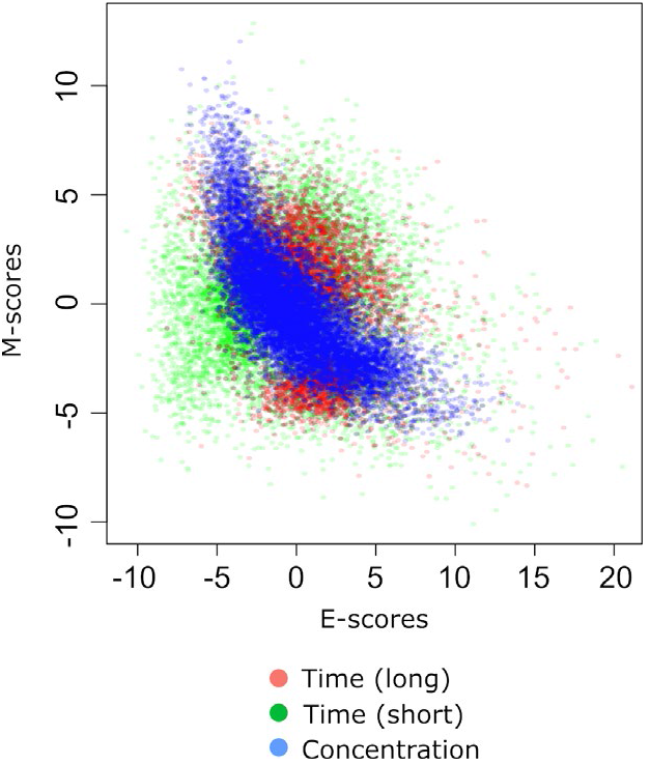
Overlap of unintegrated E- and M-scores across time and dose data. Scatter plot of E-scores (x-axis) and M-scores (y-axis) of samples from the Days 0,4,8 of time experiment (red), Days 0,1,2,3 of time experiment (green) and dose experiment (blue) prior to full integration. Note that time samples have been integrated relative to one another as in Deshmukh et al.[13], but have not been integrated with the samples from the dose experiment.

**Figure S5.**
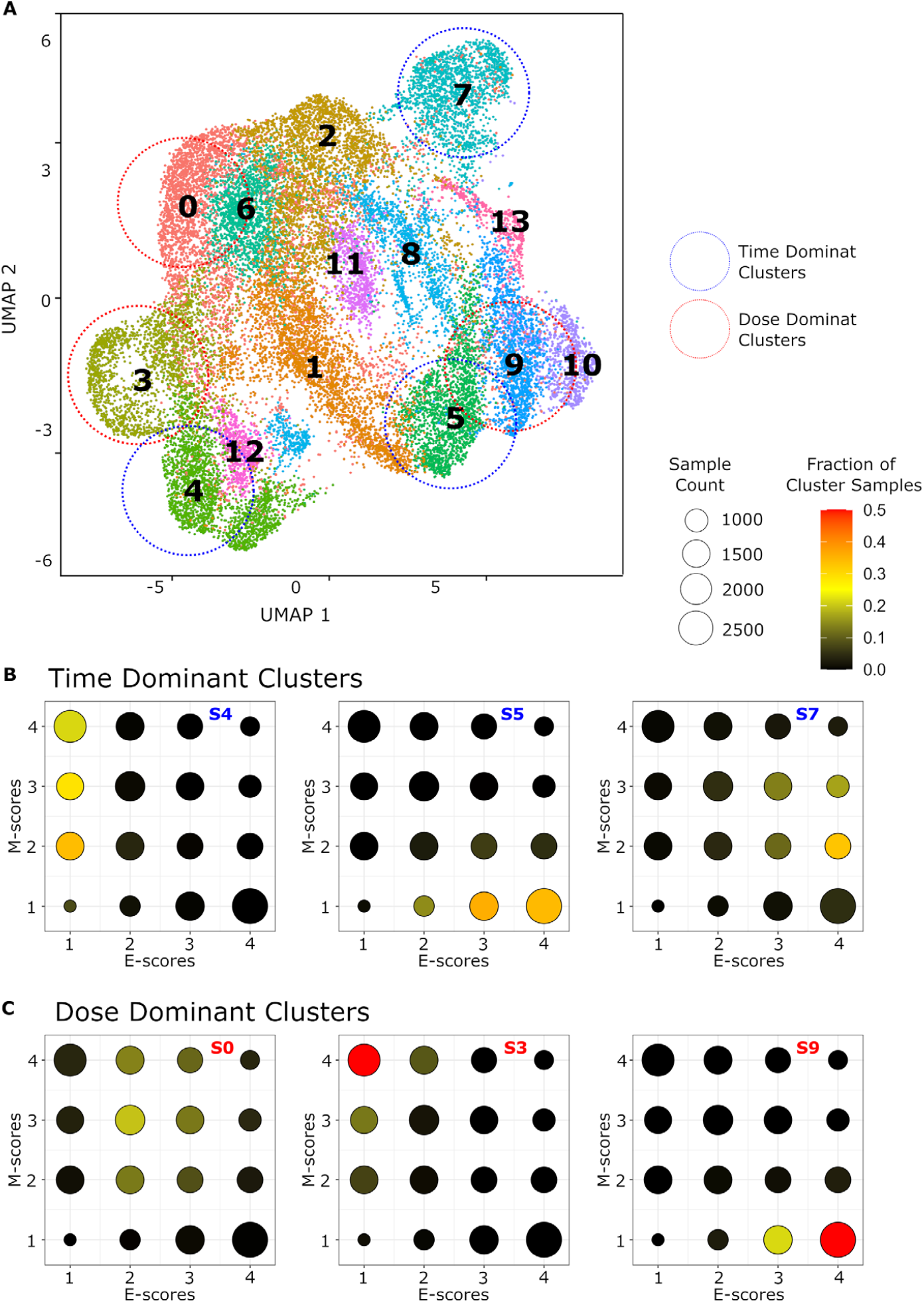
Clustering based on shared-nearest-neighbor graphs in integrated UMAP and E- and M-score space. **(A)** Projection of dose treatment single cell expression data using UMAP. The color of individual points indicates the cluster based on shared-nearest-neighbor graphs to which the sample was assigned. Dotted circles highlight three time-dominant (blue) and dose-dominant (red) clusters which we mapped to E- and M-score space. **(B)** Bubble charts showing the fraction of samples from three time dominant Nearest Neighbor clusters (4, 5, and 7 from left to right) in different segments of E- and M-score space. Each point represents a segment of E- and M-score space defined by a particular quartile of E-score (x-axis) and M-score (y-axis). The size of the point corresponds to the total number samples in the segment and the color of the point represent the fraction of samples from the corresponding cluster from 0 (black) to 0.5 (red). **(C)** Bubble charts showing the fraction of samples from three time-dominant clusters (0, 3, and 9 from left to right) in different segments of E- and M-score space. Point positions, size and color as are defined as in (B).

**Figure S6.**
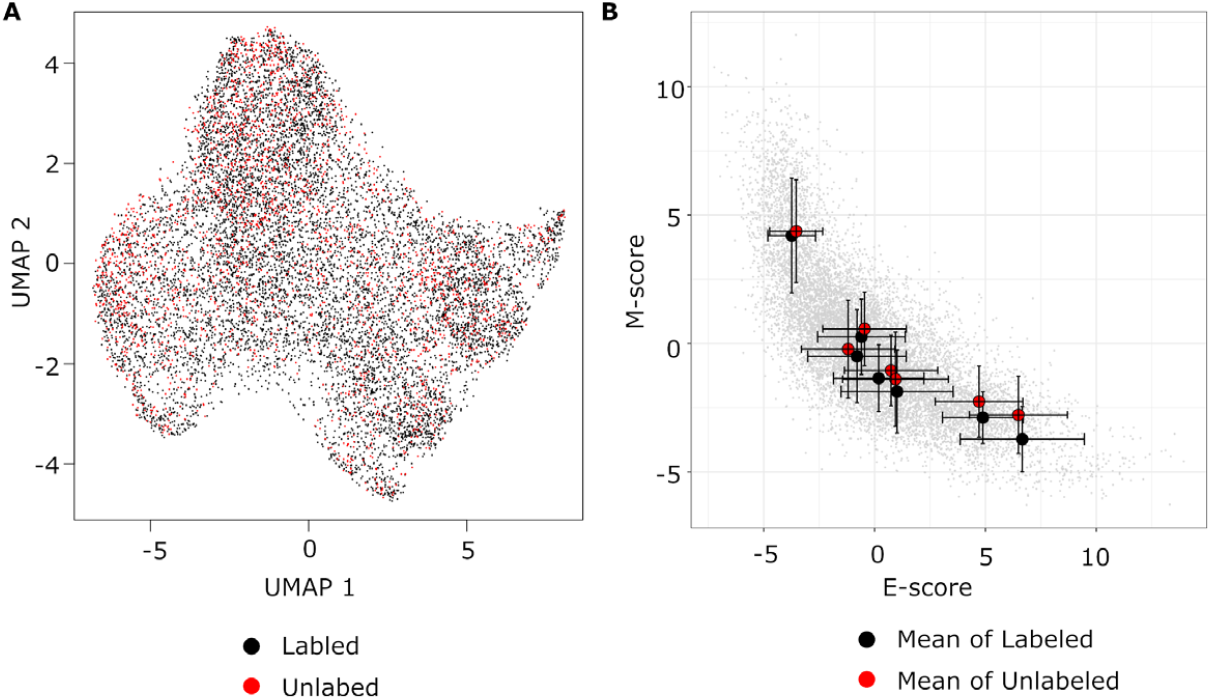
Positioning of unlabeled dose samples in UMAP space. **(A)** Projection of dose-dependent single cell expression data using UMAP. The color of individual points indicates whether the sample has a treatment level label (black) or is missing a label (red). **(B)** Scatter plots of gene set scores of E (x-axis) and M (y-axis) genes using nnPCA for dose data. Circles indicate the mean E- and M-score of samples from each shared nearest neighbor cluster, with the red dots representing labeled samples only and the blue points representing unlabeled samples. The associated error bars show the standard deviation

## References

1. Mani Sa, Guo W, Liao M-J, Eaton EN, Ayyanan A, Zhou AY, Brooks M, Reinhard F, Zhang CC, Shipitsin M, et al: The epithelial-mesenchymal transition generates cells with properties of stem cells. Cell 2008, 133:704–715.

2. Thiery JP, Acloque H, Huang RYJ, Nieto MA: Epithelial-mesenchymal transitions in development and disease. Cell 2009, 139:871–890.

3. Karacosta LG, Anchang B, Ignatiadis N, Kimmey SC, Benson JA, Shrager JB, Tibshirani R, Bendall SC, Plevritis SK: Mapping lung cancer epithelial-mesenchymal transition states and trajectories with single-cell resolution. Nat Commun 2019, 10:5587.

4. Pastushenko I, Blanpain C: EMT Transition States during Tumor Progression and Metastasis. Trends Cell Biol 2018.

5. Pastushenko I, Brisebarre A, Sifrim A, Fioramonti M, Revenco T, Boumahdi S, Van Keymeulen A, Brown D, Moers V, Lemaire S: Identification of the tumour transition states occurring during EMT. Nature 2018, 556:463–468.

6. Kröger C, Afeyan A, Mraz J, Eaton EN, Reinhardt F, Khodor YL, Thiru P, Bierie B, Ye X, Burge CB: Acquisition of a hybrid E/M state is essential for tumorigenicity of basal breast cancer cells. Proc Natl Acad Sci U S A 2019:201812876.

7. Grosse-Wilde A, Fouquier d’Herouel A, McIntosh E, Ertaylan G, Skupin A, Kuestner RE, del Sol A, Walters KA, Huang S: Stemness of the hybrid Epithelial/Mesenchymal State in Breast Cancer and Its Association with Poor Survival. PLoS One 2015, 10:e0126522.

8. Zhang J, Tian XJ, Zhang H, Teng Y, Li R, Bai F, Elankumaran S, Xing J: TGF-β -induced epithelial-to-mesenchymal transition proceeds through stepwise activation of multiple feedback loops. Sci Signal 2014, 7:ra91–ra91.

9. Wang X, Thiery JP: Harnessing Carcinoma Cell Plasticity Mediated by TGF-β Signaling. Cancers (Basel) 2021, 13:3397.

10. Hong T, Watanabe K, Ta CH, Villarreal-Ponce A, Nie Q, Dai X: An Ovol2-Zeb1 Mutual Inhibitory Circuit Governs Bidirectional and Multi-step Transition between Epithelial and Mesenchymal States. PLoS Comput Biol 2015, 11:e1004569.

11. Huang B, Lu M, Jia D, Ben-Jacob E, Levine H, Onuchic JN: Interrogating the topological robustness of gene regulatory circuits by randomization. PLoS computational biology 2017, 13:e1005456.

12. Subbalakshmi AR, Sahoo S, Biswas K, Jolly MK: A computational systems biology approach identifies SLUG as a mediator of partial Epithelial-Mesenchymal Transition (EMT). Cells Tissues Organs 2022, 211:105–118.

13. Deshmukh AP, Vasaikar SV, Tomczak K, Tripathi S, Den Hollander P, Arslan E, Chakraborty P, Soundararajan R, Jolly MK, Rai K: Identification of EMT signaling cross-talk and gene regulatory networks by single-cell RNA sequencing. Proc Natl Acad Sci U S A 2021, 118.

14. Cook DP, Vanderhyden BC: Context specificity of the EMT transcriptional response. Nat Commun 2020, 11:1–9.

15. Ramirez D, Kohar V, Lu M: Toward modeling context-specific EMT regulatory networks using temporal single cell RNA-Seq data. Frontiers in molecular biosciences 2020, 7:54.

16. Panchy N, Watanabe K, Hong T: Interpretable, Scalable, and Transferrable Functional Projection of Large-Scale Transcriptome Data Using Constrained Matrix Decomposition. Frontiers in Genetics 2021, 12:1555.

17. Tan TZ, Miow QH, Miki Y, Noda T, Mori S, Huang RYJ, Thiery JP: Epithelial-mesenchymal transition spectrum quantification and its efficacy in deciphering survival and drug responses of cancer patients. EMBO Mol Med 2014, 6:1279–1293.

18. Lee K-Y, Bae S-C: TGF-β-dependent cell growth arrest and apoptosis. BMB Reports 2002, 35:47–53.

19. Watanabe K, Panchy N, Noguchi S, Suzuki H, Hong T: Combinatorial perturbation analysis reveals divergent regulations of mesenchymal genes during epithelial-to-mesenchymal transition. npj Syst Biol Appl 2019, 5:21.

20. Freedberg IM, Tomic-Canic M, Komine M, Blumenberg M: Keratins and the keratinocyte activation cycle. J Invest Dermatol 2001, 116:633–640.

21. Hirano T, Shinsato Y, Tanabe K, Higa N, Kamil M, Kawahara K, Yamamoto M, Minami K, Shimokawa M, Arigami T: FARP1 boosts CDC42 activity from integrin αvβ5 signaling and correlates with poor prognosis of advanced gastric cancer. Oncogenesis 2020, 9:1–14.

22. Kang YH, Ji NY, Han SR, Lee CI, Kim JW, Yeom YI, Kim YH, Chun HK, Kim JW, Chung JW: ESM-1 regulates cell growth and metastatic process through activation of NF-κB in colorectal cancer. Cell Signal 2012, 24:1940–1949.

23. Wang W, Poe D, Yang Y, Hyatt T, Xing J: Epithelial-to-mesenchymal transition proceeds through directional destabilization of multidimensional attractor. Elife 2022, 11:e74866.

24. Panchy N, Azeredo-Tseng C, Luo M, Randall N, Hong T: Integrative transcriptomic analysis reveals a multiphasic epithelial–mesenchymal spectrum in cancer and non-tumorigenic cells. Front Oncol 2020, 9:1479.

25. Font-Clos F, Zapperi S, La Porta CAM: Topography of epithelial–mesenchymal plasticity. Proc Natl Acad Sci U S A 2018, 115:5902.

26. Lu M, Jolly MK, Levine H, Onuchic JN, Ben-Jacob E: MicroRNA-based regulation of epithelial-hybrid-mesenchymal fate determination. Proc Natl Acad Sci U S A 2013.

27. Jolly MK, Preca B-T, Tripathi SC, Jia D, George JT, Hanash SM, Brabletz T, Stemmler MP, Maurer J, Levine H: Interconnected feedback loops among ESRP1, HAS2, and CD44 regulate epithelial-mesenchymal plasticity in cancer. APL Bioengineering 2018, 2:031908.

28. Tian X-J, Zhang H, Xing J: Coupled reversible and irreversible bistable switches underlying TGFβ-induced epithelial to mesenchymal transition. Biophys J 2013, 105:1079–1089.

29. Nordick B, Hong T: Identification, visualization, statistical analysis and mathematical modeling of high-feedback loops in gene regulatory networks. BMC Bioinformatics 2021, 22:1–21.

30. Nordick B, Yu PY, Liao G, Hong T: Nonmodular oscillator and switch based on RNA decay drive regeneration of multimodal gene expression. Nucleic Acids Res 2022, 50:3693.

31. Dixit A, Parnas O, Li B, Chen J, Fulco CP, Jerby-Arnon L, Marjanovic ND, Dionne D, Burks T, Raychowdhury R: Perturb-Seq: dissecting molecular circuits with scalable single-cell RNA profiling of pooled genetic screens. Cell 2016, 167:1853–1866.

32. Zheng GXY, Terry JM, Belgrader P, Ryvkin P, Bent ZW, Wilson R, Ziraldo SB, Wheeler TD, McDermott GP, Zhu J: Massively parallel digital transcriptional profiling of single cells. Nat Commun 2017, 8:1–12.

33. Hao Y, Hao S, Andersen-Nissen E, Mauck Iii WM, Zheng S, Butler A, Lee MJ, Wilk AJ, Darby C, Zager M: Integrated analysis of multimodal single-cell data. Cell 2021, 184:3573–3587.

34. Oh S, Abdelnabi J, Al-Dulaimi R, Aggarwal A, Ramos M, Davis S, Riester M, Waldron L: HGNChelper: identification and correction of invalid gene symbols for human and mouse. F1000Research 2020, 9.

35. Hänzelmann S, Castelo R, Guinney J: GSVA: gene set variation analysis for microarray and RNA-Seq data. BMC Bioinformatics 2013, 14:7.

36. Sigg CD, Buhmann JM: Expectation-maximization for sparse and non-negative PCA. In Proceedings of the 25th international conference on Machine learning; 2008. 2008: 960–967.

37. Scrucca L, Fop M, Murphy TB, Raftery AE: mclust 5: clustering, classification and density estimation using Gaussian finite mixture models. The R journal 2016, 8:289.

38. Biton A: MineICA: Analysis of an ICA decomposition obtained on genomics data. R package version 1.8.0. ; 2012.

39. Wu T, Hu E, Xu S, Chen M, Guo P, Dai Z, Feng T, Zhou L, Tang W, Zhan L: clusterProfiler 4.0: A universal enrichment tool for interpreting omics data. The Innovation 2021, 2:100141.

40. Colaprico A, Silva TC, Olsen C, Garofano L, Cava C, Garolini D, Sabedot TS, Malta TM, Pagnotta SM, Castiglioni I: TCGAbiolinks: an R/Bioconductor package for integrative analysis of TCGA data. Nucleic Acids Res 2016, 44:e71–e71.

